# A novel high-throughput assay using mixed genomic DNA for fast screening germline pathogenic variants in breast cancer susceptibility genes

**DOI:** 10.1101/2021.12.23.474057

**Authors:** Qiting Wan, Li Hu, Lu Yao, Jiuan Chen, Jie Sun, Juan Zhang, Ye Xu, Yuntao Xie

**Author notes:** Qiting Wan, Li Hu and Lu Yao contributed equally to this study. **Corresponding author**: Yuntao Xie, MD, PhD, Familial & Hereditary Cancer Center, Peking University Cancer Hospital & Institute, Beijing, 100142, P. R. China.

## Abstract

The demand for genetic testing for breast cancer susceptibility genes is increasing for both breast cancer patients and healthy individuals. Here we established a novel high-throughput assay to detect germline pathogenic variants in breast cancer susceptibility genes. In general, up 10 to 50 individual genomic DNA samples were mixed together to create a mixed DNA sample and the mixed DNA sample was subjected to a next-generation multigene panel. Germline pathogenic variants in breast cancer susceptibility genes could be found in the mixed DNA sample; next, site-specific Sanger sequencing was performed to identify individuals who carried he pathogenic variant in the mixed samples. We found that the recall and precision rates were 89.9% and 92.9% when twenty individual genomic samples were mixed. Therefore, our new assay can increase an approximately 20-fold of efficacy to identify the pathogenic variants in breast cancer susceptibility genes in individuals when compared with current assay.

---

Breast cancer is the most common malignant disease in women, with up to 2.6 million new cases per year worldwide to date ^1^. Studies have reported that nearly 10% of unselected breast cancer patients carry germline pathogenic variants in breast cancer susceptibility genes ^2,3^. Among these, germline pathogenic variants in *BRCA1/2* genes accounted for 5-6% of unselected breast cancer patients, indicating that approximately one-twentieth of breast cancer patients may carry a *BRCA1* or *BRCA2* pathogenic variant ^2,3^. In addition, healthy women who carry a germline pathogenic variant of *BRCA1/2* genes are associated with a strongly increased risk of breast and ovarian cancer ^4^. Therefore, identifying germline variants in breast cancer susceptibility genes plays an important role in both healthy individuals and patients in primary prevention and optimal treatment strategies. For instance, prophylactic mastectomy could reduce the risk of breast cancer by more than ninety percent among healthy *BRCA1/2* mutation carriers ^5^. HER2-negative primary operable breast cancer patients who carry *BRCA1/2* germline pathogenic variants may benefit from adjuvant olaparib treatment based on the data of a recent OlympiA trial ^6^. As many HER2-negative breast cancer patients with *BRCA1/2* pathogenic variants may benefit from poly (ADP-ribose) polymerase (PARP) inhibitors, the potential demand for *BRCA1/2* gene testing for such patients is enormous.

It has been documented that the prevalence of *BRCA1/2* germline mutations in general populations ranges from 0.2% to 0.6% based on different ethnicities ^7–12^. The prevalence of *BRCA1/2* mutations in the Chinese Han population was 0.38% ^12^. Hence, there were approximately 5.1 million individuals carrying *BRCA1/2* mutations among the 1.3 billion Chinese Han persons ^12^.

To date, screening pathogenic variants in *BRCA1/2* or other genes (i.e., *TP53*, *PALB2*) in breast cancer patients or healthy individuals are widely applied using next-generation sequencing assays through a multigene panel; this assay usually uses an individual genomic DNA sample derived from a blood sample for genetic testing. It is cost- and time-consuming when determining germline pathogenic variants in a given gene in a huge sample of breast cancer patients or an entire general population.

In this study, we established a novel assay to meet this challenge. We combined 10 or 20 individual genomic DNA samples to be one DNA sample (mixed DNA sample), and we detected the germline variants in the mixed sample through a multigene panel and further combined them with site-specific Sanger sequencing to identify the variant in the individual. Herein, we describe this novel assay.

## Results

### General procedure of this method

In general, we had 5 mixed genomic DNA samples (see Methods section) containing 10 to up to 50 individual genomic DNA samples (each sample with equal amounts) with a total DNA amount of 1.0μg. The mixed genomic DNA samples were subjected to next-generation panel gene assays with deep sequencing (i.e., 2000x). In addition, the genomic DNA sample of each individual was also subjected to the same panel to determine the germline variants in the individual. Comparing the results of the panel sequencing of mixed DNA samples and individual DNA samples, we were able to determine that the maximum number of individual DNA samples to be mixed together could achieve optimal precision and recall rates to detect germline variants in the mixed samples. Next, when the pathogenic variants in given genes (here, we mainly focused on breast cancer susceptibility genes) were found in the mixed samples, we then screened each individual DNA sample in which they were involved in the mixed DNA sample using site-specific Sanger sequencing. Therefore, we can identify the variant in exact individuals (see the workflow in Fig.1).

**Fig. 1.**
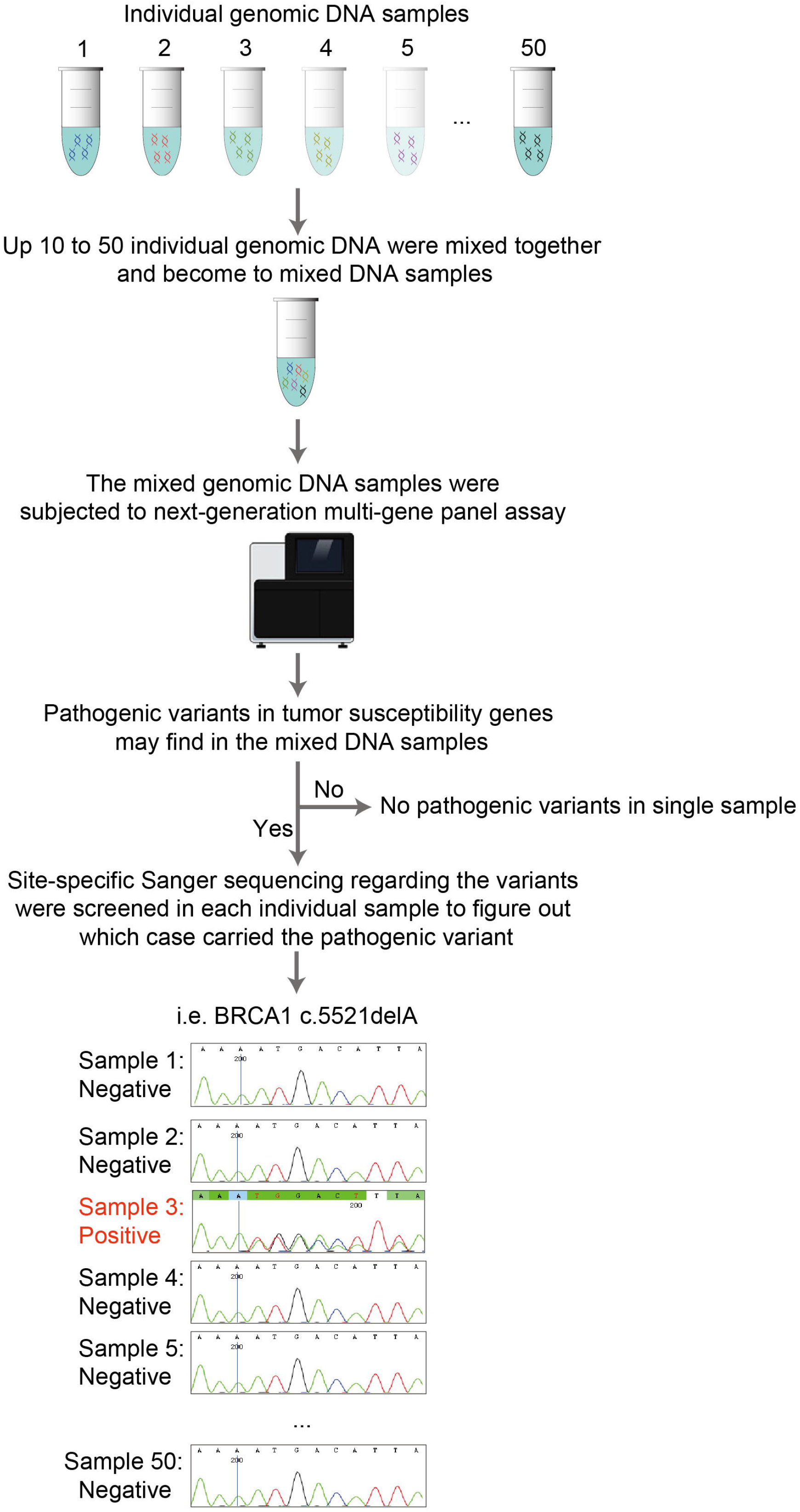
Flow diagram of the high-throughput assay using mixed genomic DNA for screening germline pathogenic variants.

### Determining the optimal number of individual DNA samples mixed together and the variants in the established set

In the established set, mixed DNA samples contained 10 to up to 50 individual DNA samples. To determine the optimal number of individual DNA samples mixed together, we calculated the precision and recall rates represented with the F1 score by comparing the results of panel sequencing of mixed DNA samples with the sequencing results of each individual DNA sample under different filtering conditions (Extended Data Fig.1-5). Consequently, the F1 scores of mixed DNA samples decreased when the number of individual DNA samples increased in the mixed samples (Fig.2A). When 10 individual DNA samples were mixed together (R10), the F1 score reached 98.2% (recall rate 98.5% and precision rate 98.0%) (Fig.2A); when 20 individual DNA samples were mixed (R20), the F1 score reached 91.3% (recall rate 89.9% and precision rate 92.9%) (Fig.2A). However, when 30 individual DNA samples were mixed, the F1 score was 87.0%, indicating that more than 10% of variants might be lost (Fig.2A and Extended Data Fig.3). Therefore, the optimal number of individual DNA samples mixed together was twenty, with less than 10.0% of variants being lost; when the number of individual DNA samples was restricted to ten, less than 2.0% of variants might be lost (Fig.2A).

**Fig. 2.**
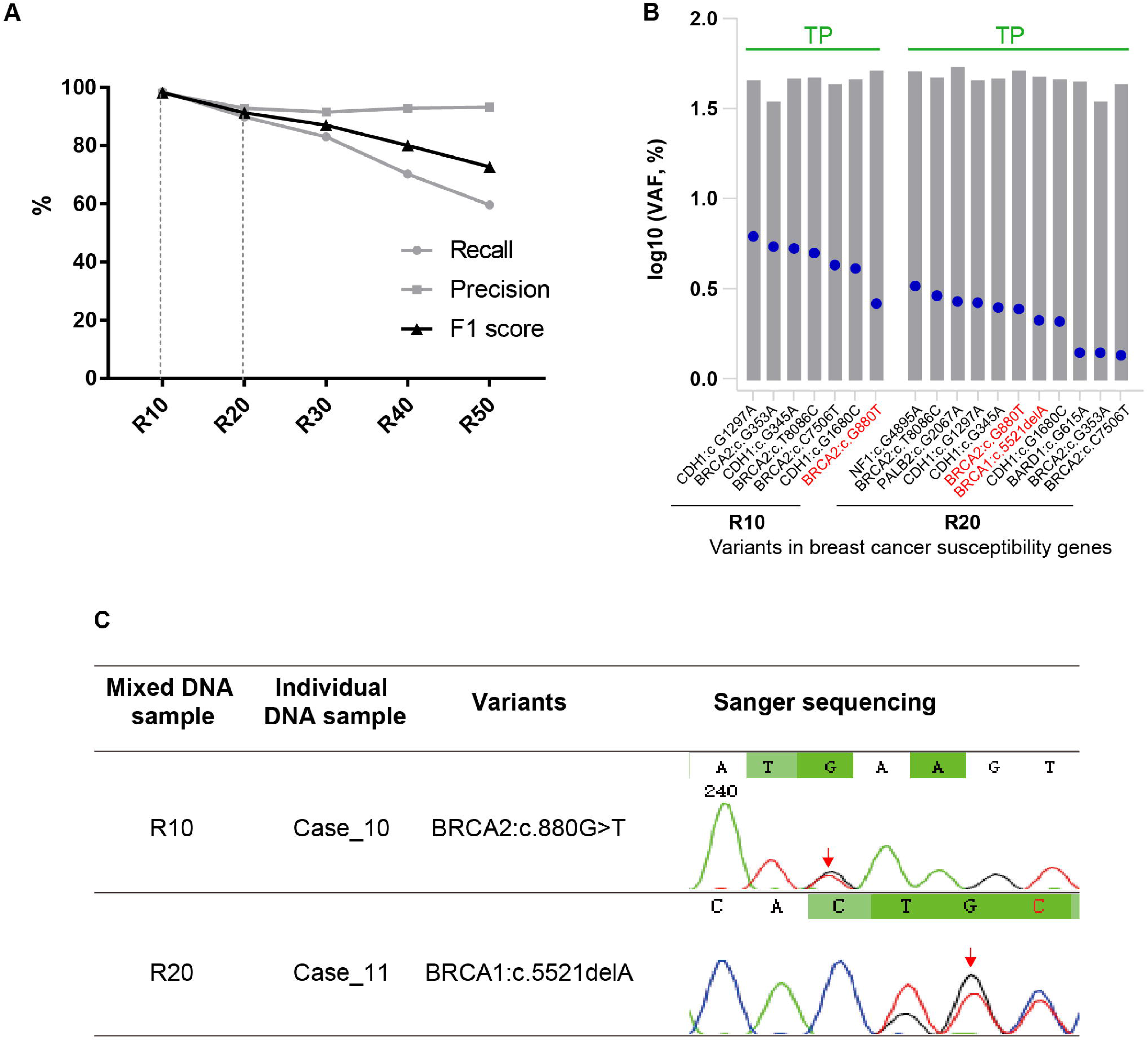
To establish the optimal number of individual DNA samples mixed and determine the variants. The mixed genomic DNA samples and each individual DNA sample were subjected to next generation 560-genes panel assay, comparing the results of the panel sequencing of mixed DNA samples and individual DNA samples, we were able to establish a maximum number of individual DNA sample to be mixed and to figure out the variants in the mixed samples. **A.** Recall and precision rates at the optimal variant filtering condition for mixed genomic DNA sequencing. Variants from the individual sample sequencing were filtered at a depth >50x, and the variant allele frequency > 3.0%. Variants from the 5 mixed genomic DNA (from R10 to R50) sequencing were filtered at the depth of variant-supporting bases on forward and reverse strand >1x, the variant allele frequency >2.5% for R10, >1.0% for R20, and >0.5% for R30 to R50, and P value from Fishers exact test < 1e −3 for R10, < 1e-2 for R20 to R50. **B.** Variants in twelve breast cancer susceptibility genes were detected in mixed DNA samples R10 and R20. We calculated the log10 (VAF, %) values for variants detected in the individual sample (gray bars) and the mixed DNA samples (blue dots). Variants present in both individual DNA samples and mixed DNA samples (R10 to R20) were true positive (TP). Pathogenic variants were marked with red. **C.** Individuals carrying the pathogenic variants in breast cancer susceptibility genes were identified by site-specific Sanger sequencing.

We then focused on variants in breast cancer susceptibility genes. All of the pathogenic variants in the twelve breast cancer susceptibility genes were successfully detected in the mixed DNA samples (R10 and R20) when compared with the panel sequencing data of the 20 individual DNA samples (Fig.2B). In mixed DNA sample R10, one *BRCA2* pathogenic variant c.880G>T was found (Table 1 and Fig.2B). To identify which individual carried the pathogenic variant in R10 (a total of 10 individual DNA samples from case 1 to case 10), site-specific Sanger sequencing regarding this variant was performed in the ten samples. We found that case 10 contained these variants (Table 2 and Fig.2C). Similarly, two pathogenic variants (*BRCA2* c.880G>T and *BRCA1* c.5521delA) were found in mixed sample R20 (a total of 20 individual DNA samples from case 1 to case 20) (Table 1 and Fig.2B). We found that case 10 contained *BRCA2* c.880G>T and case 11 contained *BRCA1* by using site-specific Sanger sequencing regarding the two variants (Table 2 and Fig. 2C). Using this procedure, we can also identify the pathogenic variants in other cancer susceptibility genes (data not shown).

**Table 1.** Pathogenic variants in breast cancer susceptibility genes detected in mixed DNA samples by 550-panel sequencing.

| **Mixed sample** | **BRCA1** | **BRCA2** | **PALB2** | **ATM** | **CHEK2** | **BARD1** | **RAD51C** | **RAD51D** | **TP53** | **CDH1** | **PTEN** | **NF1** |
| --- | --- | --- | --- | --- | --- | --- | --- | --- | --- | --- | --- | --- |
| R10 | - | c.880G>T | - | - | - | - | - | - | - | - | - | - |
| R20 | c.5521delA | c.880G>T | - | - | - | - | - | - | - | - | - | - |
| R30 | c.5521delA,<br>c.3329dupA | - | c.751C>T | - | - | - | - | - | - | - | - | - |
| R40 | c.5521delA | - | c.751C>T | - | - | - | - | - | - | - | - | - |
| R50 | c.5521delA,<br>c.3329dupA | c.880G>T | c.751C>T | - | - | - | - | - | - | - | - | - |

**Table 2.** To identify the variant carriers found in the mixed DNA samples by site-specific Sanger sequencing.

| Mixed DNA sample | Individual DNA sample | BRCA2: c.880G>T | BRCA1:c.5521delA |
| --- | --- | --- | --- |
| R10 | Case_1 | × | - |
|  | Case_2 | × | - |
|  | Case_3 | × | - |
|  | Case_4 | × | - |
|  | Case_5 | × | - |
|  | Case_6 | × | - |
|  | Case_7 | × | - |
|  | Case_8 | × | - |
|  | Case_9 | × | - |
|  | Case_10 | ○ | - |
| R20 | Case_1 | × | × |
|  | Case_2 | × | × |
|  | Case_3 | × | × |
|  | Case_4 | × | × |
|  | Case_5 | × | × |
|  | Case_6 | × | × |
|  | Case_7 | × | × |
|  | Case_8 | × | × |
|  | Case_9 | × | × |
|  | Case_10 | ○ | × |
|  | Case_11 | × | ○ |
|  | Case_12 | × | × |
|  | Case_13 | × | × |
|  | Case_14 | × | × |
|  | Case_15 | × | × |
|  | Case_16 | × | × |
|  | Case_17 | × | × |
|  | Case_18 | × | × |
|  | Case_19 | × | × |
|  | Case_20 | × | × |

Notes: “○” indicates the variant was detected in the corresponding cases; “×” indicates the variant was not detected in the corresponding cases.

### Validating the method in independent cohorts of breast cancer patients and healthy women

In the validation set, to explore whether the method was reliable, we applied this method in another two independent validation sets including twenty breast cancer patients and twenty healthy women. In the mixed DNA sample P10 (containing 10 patient DNA samples, from case 1 to case 10), the F1 score was 96.1% (recall rate 96.9% and precision rate 95.3%); in the mixed DNA sample P20 (containing 20 patient DNA samples, from case 1 to case 20), the F1 score was 87.4% (recall rate 96.0% and precision rate 80.2%) (Fig. 3A); several pathogenic variants in the 12 breast cancer susceptibility genes were found in the mixed DNA samples (Fig. 3B and Supplementary Table 4). By site-specific Sanger sequencing, we were able to identify which case carried the pathogenic variant (Supplementary Table 5 and Extended Data Fig.6).

**Fig. 3.**
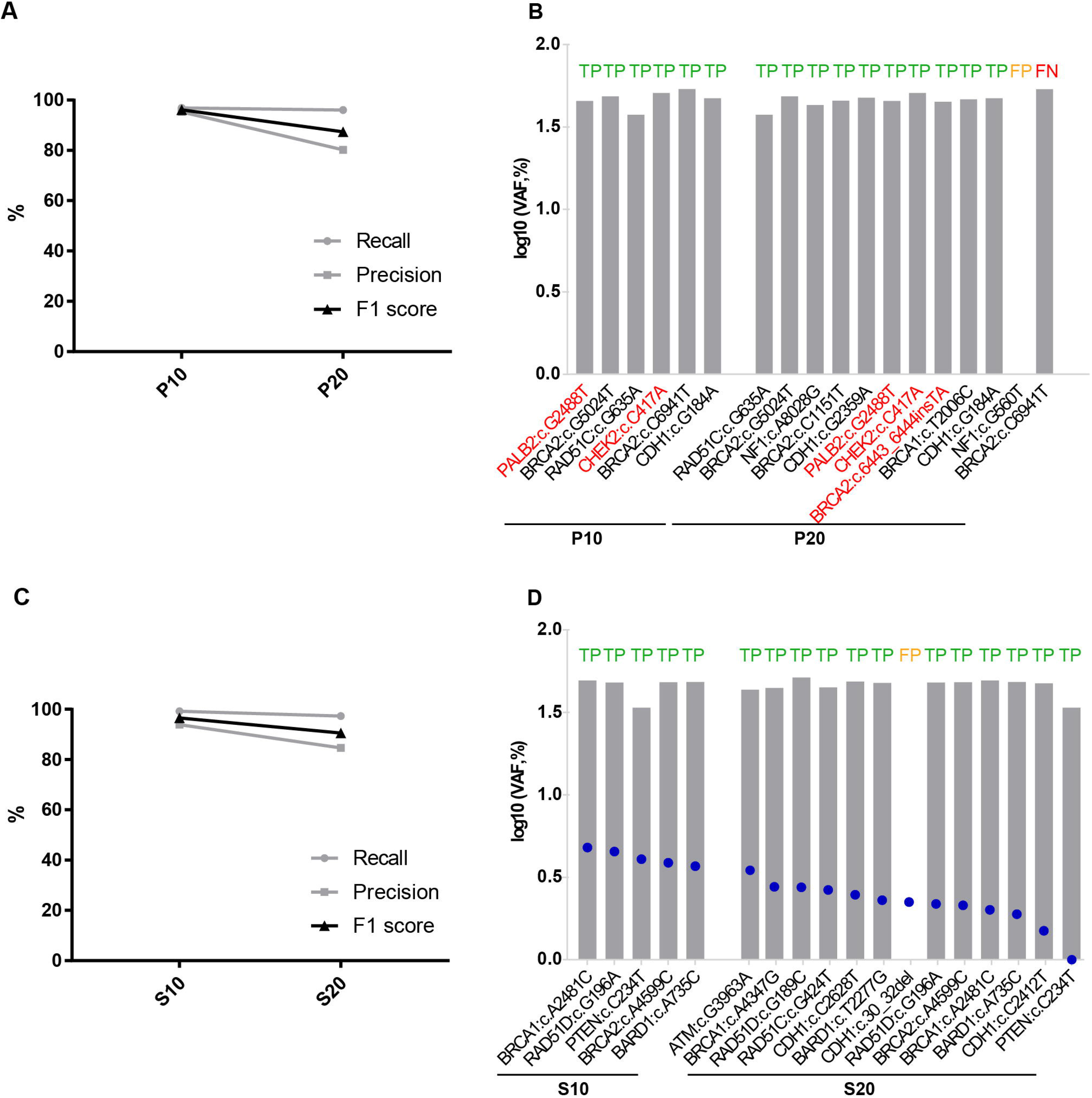
Independent validation of mixed DNA sequencing in additional breast cancer patients and healthy women. **A,C**. We performed individual DNA sample sequencing and mixed DNA sample sequencing in an additional twenty breast cancer patients (P10, ten samples mixed; P20, twenty samples mixed) and twenty healthy women (S10, ten samples mixed; S20, twenty samples mixed). **B,D**. We calculated the log10 (VAF, %) values for variants detected in individual DNA samples (gray bars) and mixed DNA samples (blue dots). Variants present in both individual DNA samples and mixed DNA samples were true-positive (TP), variants detected only in the individual DNA sample but not in the mixed DNA sample were false-negative (FN), and variants detected only in the mixed DNA sample but not in the individual DNA sample were false-positive (FP). Pathogenic variants were marked in red.

In the mixed DNA sample S10 (containing 10 healthy women DAN samples, from healthy women No. 1 to No. 10), the F1 score was 96.5% (recall rate 99.2% and precision rate 93.9%) (Fig.3C); in the mixed DNA sample S20 (containing 20 healthy women DAN samples, from healthy women No. 1 to No. 20), the F1 score was 90.5% (recall rate 97.3% and precision rate 84.6%) (Fig.3C). Since no pathogenic variants in the breast cancer susceptibility genes were found in the mixed DNA samples from the healthy women, we therefore listed the unknown variants in these genes (Fig.3D and Supplementary Table 6). Similarly, using site-specific Sanger sequencing, healthy women who carried this variant were identified (Supplementary Table 7 and Extended Fig.7).

We also noticed that less than 10% of false-negative or false-positive variants might occur in the mixed samples, especially when 20 individual DNA samples were mixed together. For example, one false-positive variant in the *NF1* gene and one false negative variant in the *BRCA2* gene were found in P20 (Fig.3B), and one false-positive variant in *CDH1* was found in S20 (Fig.3D). However, no false-negative or false-positive variants in these genes were found when ten individual DNA samples were mixed regardless of breast cancer patients or healthy women.

## Discussion

In this study, we established a novel high-throughput assay to detect germline pathogenic variants in breast cancer susceptibility genes by combining next-generation multigene panels and site-specific Sanger sequencing. In general, we combined 10 to 20 individual genomic DNA samples to create a mixed DNA sample.

This mixed DNA sample was subjected to a next-generation multigene panel (550-gene panel in this study). Germline pathogenic variants in breast cancer susceptibility genes could be found in the mixed DNA sample; next, site-specific Sanger sequencing regarding the mutation site was screened for each individual DNA sample; thus, the mutation in the exact case can be identified.

Currently, germline pathogenic variants in cancer susceptibility genes are routinely detected by next-generation multigene panels using individual genomic DNA. Compared to the current assay, our new assay not only accelerated the efficacy by 10-to 20-fold but also dramatically decreased the cost of detection. Our assay can be used to detect germline pathogenic variants in breast cancer susceptibility genes in breast cancer patients and healthy women. It is well documented that the prevalence of pathogenic variants in known breast cancer susceptibility genes is approximately 10% in unselected breast cancer patients, and *BRCA1/2* account for approximately 5.0-6.0% ^3^. If we are intended to detect *BRCA1/2* germline variants in one hundred breast cancer patients, we only needed to detect 5 mixed DNA samples (one mixed sample containing 20 individual DNA samples) instead of 100 DNA samples. The chances of the supposed five *BRCA1/2* pathogenic variants might range from one variant in each mixed sample to five variants in only one mixed sample. If a mixed sample did not contain any *BRCA1/2* pathogenic variants, the mixed sample could be ignored, suggesting that the corresponding 20 individual DNA samples were negative for *BRCA1/2* mutations; in contrast, for those mixed samples containing one to five *BRCA1/2* variants, we used site-specific Sanger sequencing regarding the mutation sites to determine the variant in the exact cases. Therefore, if we wanted to detect *BRCA1/2* mutations in ten thousand breast cancer patients, we only needed to detect five hundred mixed DNA samples.

This new assay might be used not only in breast cancer patients but also in healthy women. The prevalence of *BRCA1/2* mutations in the Chinese Han population is 0.38% ^12^. Thus, there are approximately 40 *BRCA1/2* mutations in ten thousand healthy Chinese Han women, as five hundred mixed DNA samples could cover the ten thousand healthy women individual DNA samples, and 40 *BRCA1/2* mutations could maximally occur in 40 of 500 mixed samples. Therefore, site-specific Sanger sequencing was only performed in 800 healthy women individual DNA samples to determine the 40 *BRCA1/2* mutations. This could save a huge amount of time and cost and make the strategy of screening *BRCA1/2* mutations in the entire population possible and fast. In addition, this assay was not restricted to detecting germline variants in breast cancer susceptibility genes, and it was also extended to detect other cancer susceptibility genes.

Despite the many advantages of this new assay, the disadvantage of this assay might be that it misses approximately 10% of mutations when the mixed DNA sample contains 20 individual DNA samples from breast cancer patients. Therefore, caution is required when the assay is applied in clinical practice.

To date, the demand for *BRCA1/2* detection is increasing in both breast cancer patients and healthy women; therefore, our new assay is useful for meeting these challenges.

## Methods

### Patients and healthy women

In the established set, 50 patients with operable primary breast cancer who were treated at the Breast Center of the Peking University Cancer Hospital from December 2016 to February 2017 and were unselected for age of diagnosis and family history were included. All 50 patients were Han Chinese, and the age at diagnosis ranged from 16 to 74 (median 44). Detailed information on the 50 patients is presented in Supplementary Table 1. In the validation set, 20 patients with operable primary breast cancer were randomly selected from a large cohort of 8085 breast cancer patients as described previously ^3^. The 20 patients were also Han Chinese, and the age at diagnosis ranged from 31 to 77 (median 47). The 20 healthy women resided in the Beijing area and were all Chinese Han with ages from 23 to 50 (median 40).

### Established Set

#### Sample preparation

Fifty unselected breast cancer patients (case 1 to case 50) were included in the established set. Genomic DNA was extracted from peripheral blood samples by using a blood genome DNA isolation kit (BioTeke, China). The genomic DNA from the different cases was mixed together with equal quantities of DNA content for each case (Supplementary Table 2). First, DNA samples from ten cases were mixed together; then, the number of cases was increased every ten cases. Based on this principle, ten genomic DNA samples from case 1 to case 10 (100 ng per case) were mixed together, and we called the mixed DNA sample R10 with a total of 1.0 μg DNA. Then, 20 genomic DNA samples from case 1 to case 20 (50 ng per case) were mixed together (called R20), 30 genomic DNA samples from case 1 to case 30 (33 ng per case) were mixed together (called R30), 40 genomic DNA samples from case 1 to case 40 (25 ng per case) were mixed together (called R40), and 50 genomic DNA samples from case 1 to case 50 (20 ng per case) were mixed together (called R50). Therefore, there were 5 mixed DNA samples containing 10 to 50 individual DNA samples with a total of 1. 0μg DNA for each mixed DNA sample. The 5 mixed DNA samples were subjected to next-generation sequencing assays using a 550-gene panel (see below). In addition, individual DNA samples (1.0 μg) from case 1 to case 50 were also subjected to 550-gene panel target sequencing.

#### Library preparation and target sequencing

The sequencing library was generated with the TruSeq Nano DNA HT Sample Prep Kit (Illumina USA) for both the mixed DNA samples and individual DNA samples. Target sequencing was performed for all the exons of 550 cancer-related genes (Agilent SureSelect XT Custom kit; Agilent Technologies, Palo Alto, CA, http://www.agilent.com). Twelve breast cancer susceptibility genes reported recently were included in these 550 genes^13,14^ (Supplementary Table 3). The libraries were sequenced on the NovaSeq6000 platform. Panel sequencing was performed for each mixed DNA sample at a depth of 2000-fold and 500-fold for each individual DNA sample. High-throughput sequencing was performed for each captured library independently to ensure that all samples met the desired average fold change.

#### Variant calling

For both the mixed DNA samples and individual DNA samples, VarScan2 was used to call germline single nucleotide variations (SNVs), insertions and deletions (InDels)^15^. Variant call format (VCF) was annotated by ANNOVAR^16^. Filtering conditions were set to identify germline variants for both mixed and individual samples as follows: 1) include heterozygous variants located at the exon or splicing region; 2) remove variants with mutant allele frequency (MAF) over 1% in the population databases, including the 1000 Genomes Project^17^, Exome Sequencing Project (ESP6500), and the Exome Aggregation Consortium (ExAC); and 3) remove variants located at segmental duplications.

Subsequently, variants detected in mixed and individual DNA samples were filtered under different conditions. For mixed DNA samples, variants were filtered out at a depth of variant-supporting bases on the forward and reverse strands > 1-fold; in particular, the cutoff for variant allele frequency increased stepwise from >0.5% to >3.0%, and the cutoff for the P value from Fisher’s exact test increased stepwise from <1e-10 to <1. For each individual DNA sample, variants were filtered out at a sequencing depth > 50-fold and a variant allele frequency > 30.0%.

#### Accuracy of variant detection in mixed DNA samples

To evaluate the variant detection accuracy of mixed DNA samples, we calculated the precision and recall rates under different cutoffs of variant allele frequency and P value from Fisher’s exact test when compared with the sequencing results of the individual DNA samples. The F1 score was calculated to estimate the variant detection accuracy when different numbers of samples were mixed. In addition, the optimal number of samples to be mixed as well as the proper cutoffs for variant allele frequency and P value were further confirmed in the validation set.

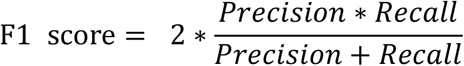

#### To identify pathogenic variants in individual cases via site-specific Sanger sequencing

We mainly focused on the twelve breast cancer susceptibility genes in the 550-gene panel. Pathogenic variants in these susceptibility genes might be found in the five mixed DNA samples (from R10 to R50). For example, pathogenic variants were found in the mixed DNA sample R10 (Table 1), which contained the individual DNA samples from cases 1 to 10. We then screened the individual DNA sample from case 1 to case 10 by site-specific Sanger sequencing regarding the pathogenic variants (Table 2). Therefore, we were able to identify the variants that occurred in each case.

### Validation set

To validate the precision and recall rates of the method, we constructed a validation set that contained an additional twenty breast cancer patients and twenty healthy individuals. For the twenty patients (from case 1 to case 20), the genomic DNA from case 1 to case 10 (100 ng per case) was mixed together, and we called the mixed DNA sample P10 with a total of 1.0μg DNA; the genomic DNA from case 1 to case 20 (50 ng per case) was mixed together (called P20). Then, the two mixed DNA samples with a total of 1μg DNA per sample were subjected to a 550-gene panel sequencing with 2000-fold. In addition, each individual DNA sample (1.0μg) from case 1 to case 20 was also subjected to 550-gene panel target sequencing with 500-fold. Similarly, regarding the twenty healthy individuals (case 1 to case 20), the genomic DNA from case 1 to case 10 (100 ng per case) was mixed together (called S10), and the genomic DNA from case 1 to case 20 (50 ng per case) was mixed together (called S20). Then, the two mixed DNA samples with a total of 1μg DNA were subjected to a 550-gene panel with 2000-fold. In addition, each healthy individual DNA sample (1. 0μg) from case 1 to case 20 was also subjected to 550-gene panel target sequencing with 500-fold. The procedures of site-specific Sanger sequencing to identify the variants in breast cancer patients or healthy individuals were described above.

## Data availability

The datasets generated and/or analyzed during the current study are not publicly available but are available from the corresponding author on reasonable request.

## Author contributions

Y.X. had full access to all the data in the study and take responsibility for the integrity of the data and the accuracy of the data analysis.

*Study concept and design*: Y.X.

*Acquisition, analysis, or interpretation of data*: Q.W., L.H., L.Y. and Y.X.

*Drafting of the manuscript*: Q.W. and Y.X.

*Critical revision of the manuscript for important intellectual content*: Y.X.

*Obtained funding*: Y.X.

*Administrative, technical, or material support*: Q.W., L.H., L.Y., J.C., J.S., J.Z., Y.X. and Y.X.

*Study supervision*: Y.X.

## Extented data

Extended data Fig.1-Fig.5

## Supplementary information

Supplementary Table 1-5

