## Extended Data for "A novel high-throughput assay using mixed genomic DNA for fast screening germline pathogenic variants in breast cancer susceptibility genes"

Extended Data Fig.1

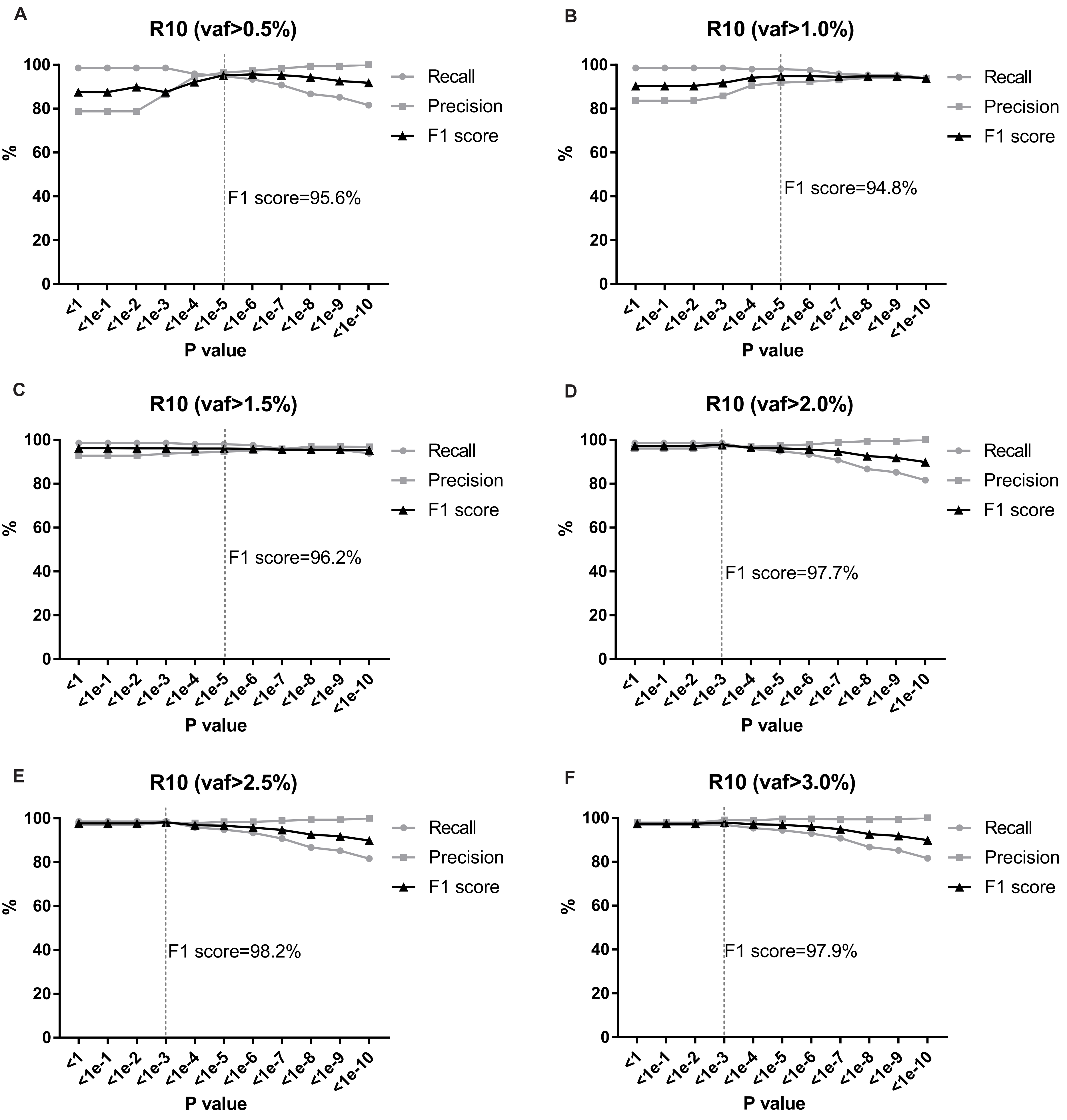

**Extended Data Fig.1 Recall and precision rates at a series of filtering condition for the genomic DNA-mixed sequencing in R10.** Variants from the single sample sequencing were filtered at the depth >50x, and the variant allele frequency >3.0%. Variants from the genomic DNA-mixed sequencing were filtered at the depth of variant-supporting bases on forward and reverse strand >1x, P-value from Fisher's exact test from <1e-10 to <1, and the variant allele frequency from >0.5% to 3.0%. Vaf, variant allele frequency.

Extended Data Fig.2

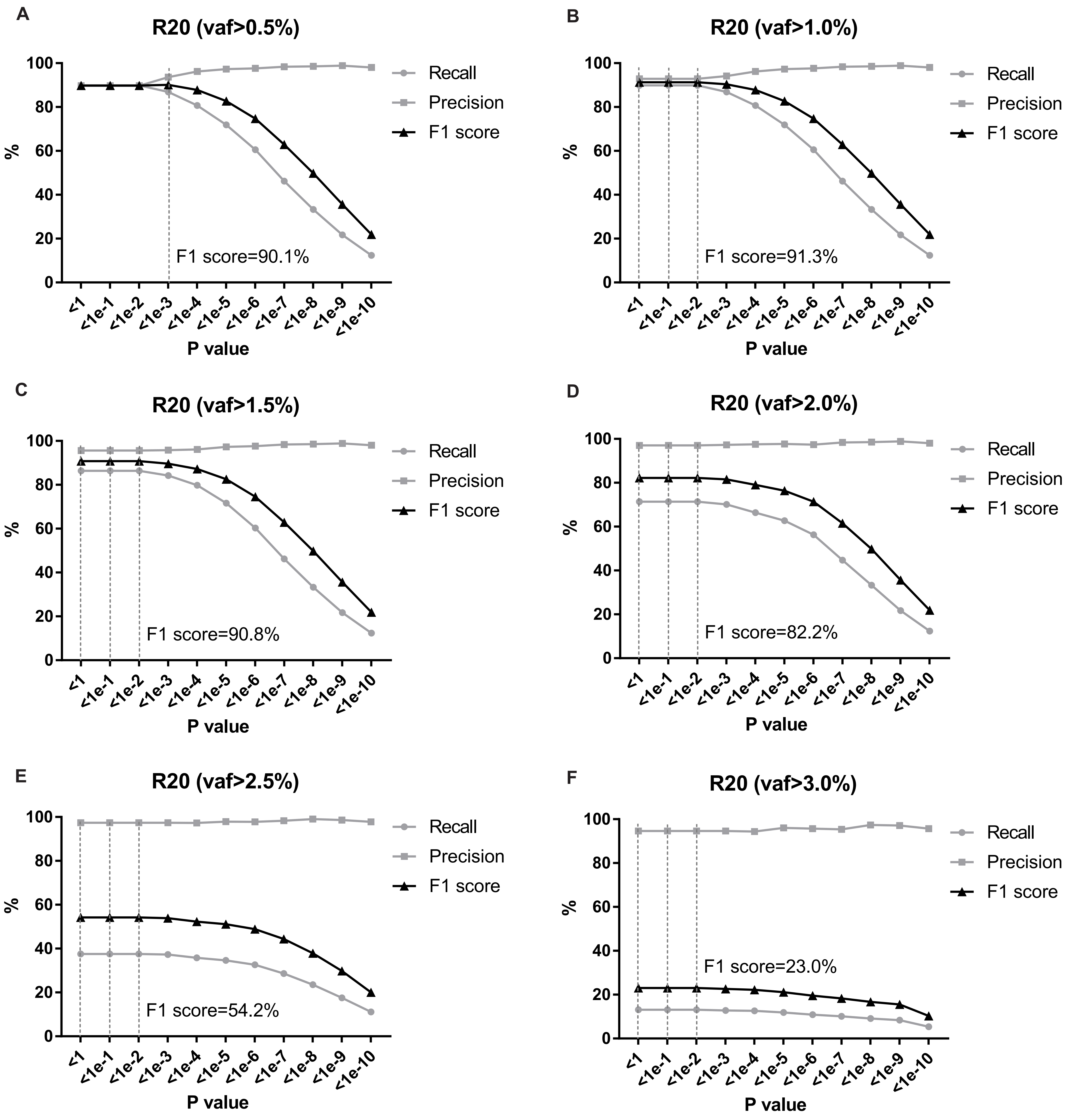

**Extended Data Fig.2 Recall and precision rates at a series of filtering condition for the genomic DNA-mixed sequencing in R20.** Variants from the single sample sequencing were filtered at the depth >50x, and the variant allele frequency >3.0%. Variants from the genomic DNA-mixed sequencing were filtered at the depth of variant-supporting bases on forward and reverse strand >1x, P-value from Fisher's exact test from <1e-10 to <1, and the variant allele frequency from >0.5% to 3.0%. Vaf, variant allele frequency.

Extended Data Fig.3

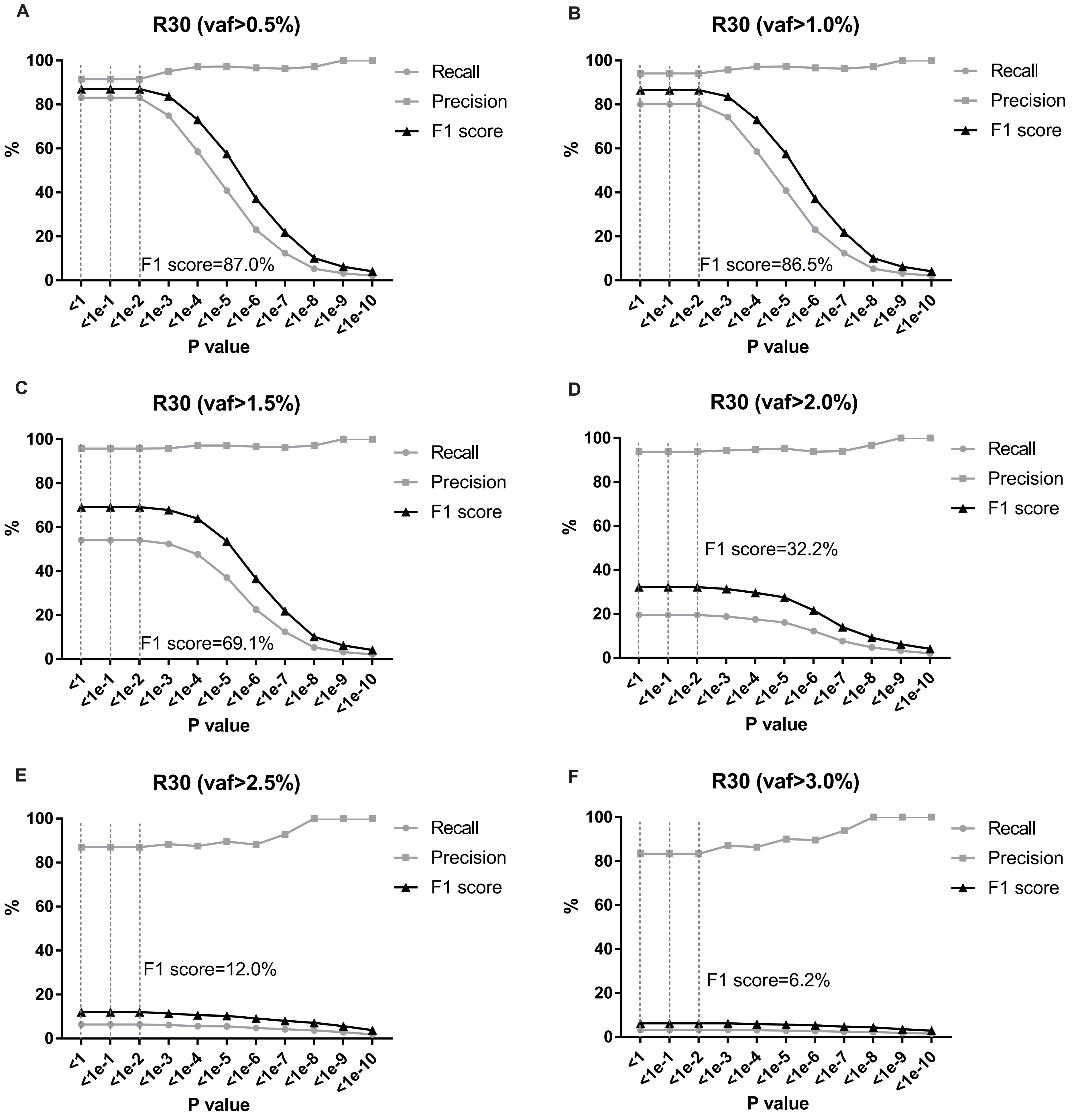

**Extended Data Fig.3 Recall and precision rates at a series of filtering condition for the genomic DNA-mixed sequencing in R30.** Variants from the single sample sequencing were filtered at the depth >50x, and the variant allele frequency >3.0%. Variants from the genomic DNA-mixed sequencing were filtered at the depth of variant-supporting bases on forward and reverse strand >1x, P-value from Fisher's exact test from <1e-10 to <1, and the variant allele frequency from >0.5% to 3.0%. Vaf, variant allele frequency.

Extended Data Fig.4

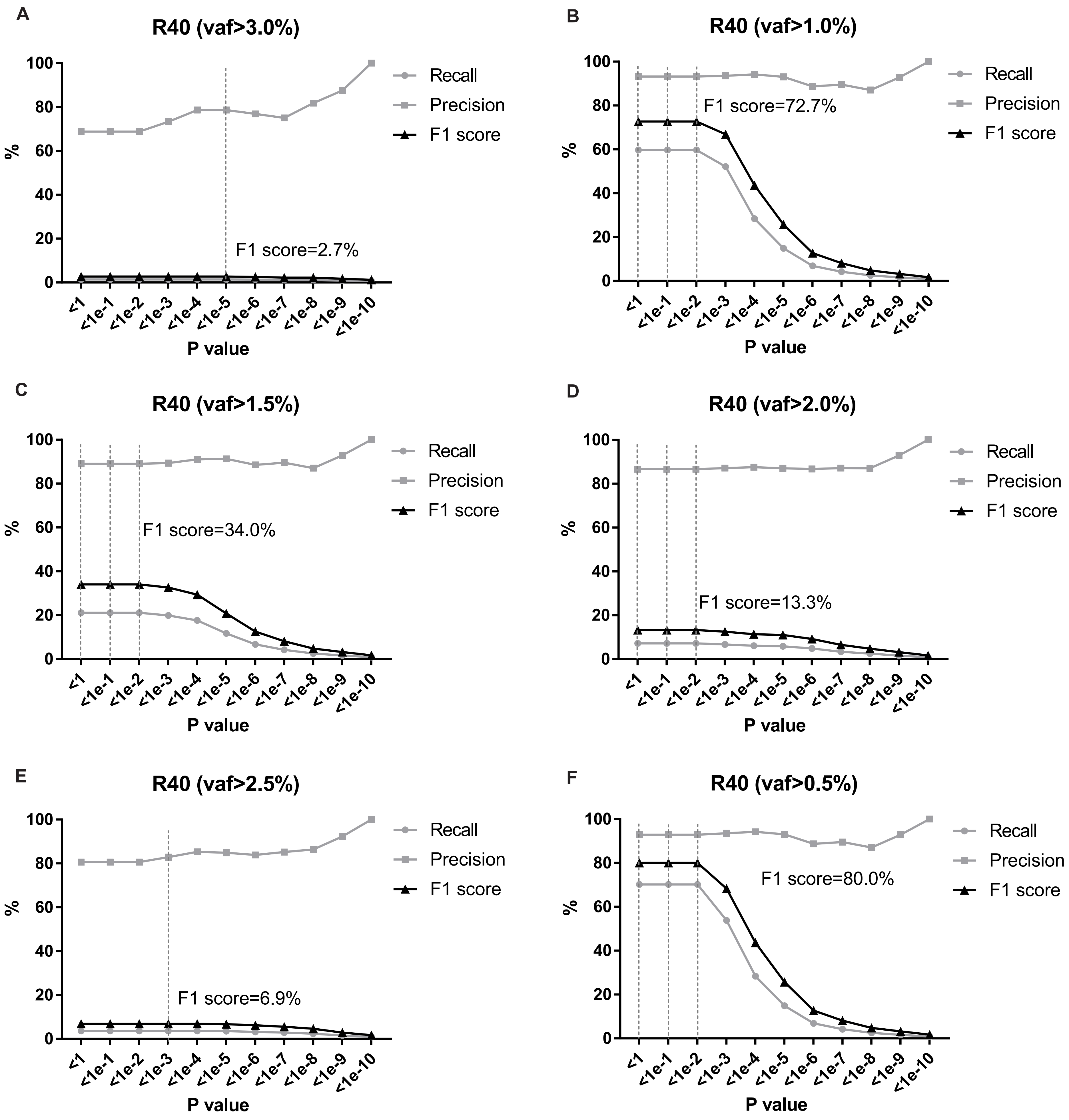

**Extended Data Fig.4 Recall and precision rates at a series of filtering condition for the genomic DNA-mixed sequencing in R40.** Variants from the single sample sequencing were filtered at the depth >50x, and the variant allele frequency >3.0%. Variants from the genomic DNA-mixed sequencing were filtered at the depth of variant-supporting bases on forward and reverse strand >1x, P-value from Fisher's exact test from <1e-10 to <1, and the variant allele frequency from >0.5% to 3.0%. Vaf, variant allele frequency.

Extended Data Fig.5

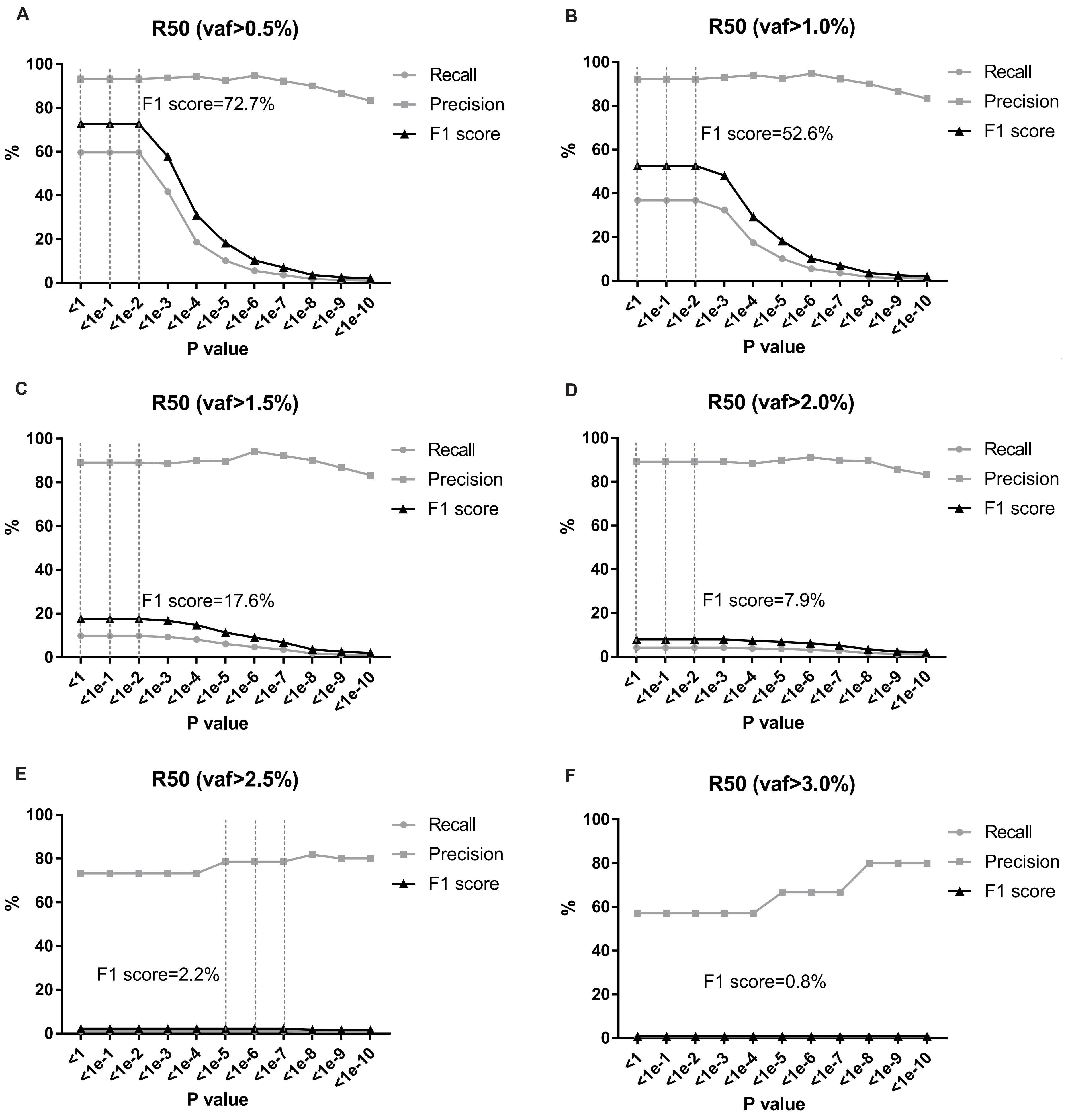

**Extended Data Fig.5 Recall and precision rates at a series of filtering condition for the genomic DNA-mixed sequencing in R50.** Variants from the single sample sequencing were filtered at the depth >50x, and the variant allele frequency >3.0%. Variants from the genomic DNA-mixed sequencing were filtered at the depth of variant-supporting bases on forward and reverse strand >1x, P-value from Fisher's exact test from <1e-10 to <1, and the variant allele frequency from >0.5% to 3.0%. Vaf, variant allele frequency.

**Extended Data Fig.6**

| Mixed DNA sample | Individual DNA sample | Variants | Sanger sequencing |
|---|---|---|---|
| P10 | Patient_2 | PALB2:c.2488G>T | 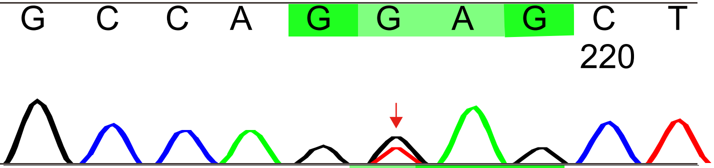 |
| P10 | Patient_2 | CHEK2:c.417C>A | 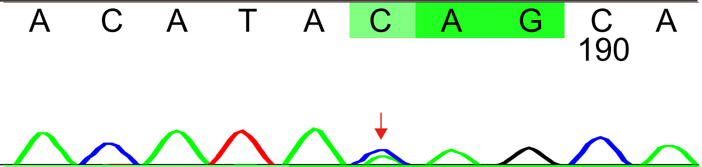 |
| P20 | Patient_12 | BRCA2: c.6443_6444insTA | 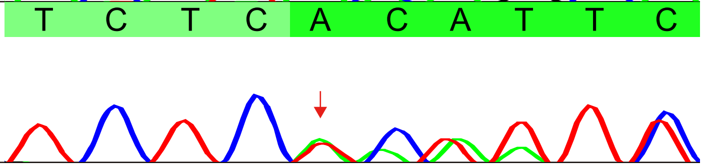 |

**Extended Data Fig.6 Individuals carrying deleterious variants in breast cancer susceptibility genes were found out by site-specific Sanger sequencing in the validation sets of patients (P10 and P20).**

### Extended Data Fig.7

| Mixed DNA sample | Individual DNA sample | Variants | Sanger sequencing |
|---|---|---|---|
| S10 | Healthy_2 | BRCA1:2481A>C | 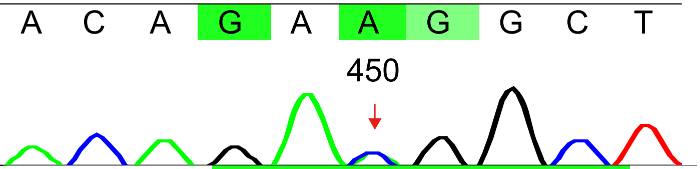  |
| S10 | Healthy_5 | BRCA2:c.4599A>C | 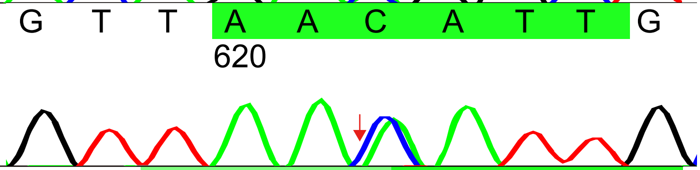  |
| S10 | Healthy_6 | RAD51D: c.196G>A | 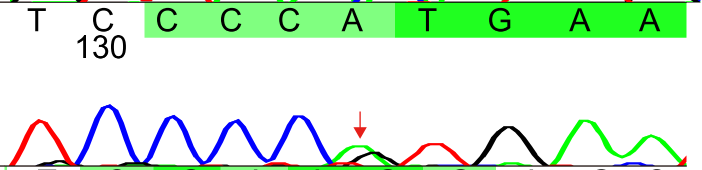  |
| S20 | Healthy_20 | BARD1: c.2277T >G | 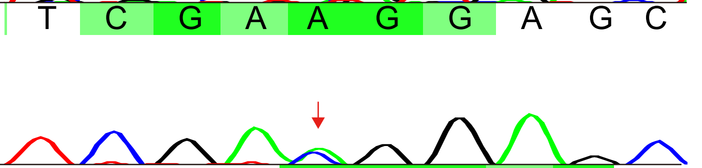  |
| S20 | Healthy_20 | RAD51C: c.424G>T | 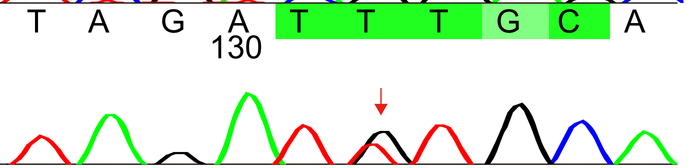  |

**Extended Data Fig.7 Individuals carrying unknown variants in breast cancer susceptibility genes were found out by site-specific Sanger sequencing in the validation sets of healthy women (S10 and S20).**
