## Supplementary Table for "A novel high-throughput assay using mixed genomic DNA for fast screening germline pathogenic variants in breast cancer susceptibility genes"

**Supplementary information**

### Supplementary Table 1. The detailed information of the fifty patients in the establish set.

| **Case ID** | **Age** | **Gender** | **Ethnics** | **Tumor location** | **Family history of BCOC** | **Tumor size, cm** | **TNM** | **Histology** | **ER** | **PR** | **HER2** | **TNBC** |
| --- | --- | --- | --- | --- | --- | --- | --- | --- | --- | --- | --- | --- |
| Case_1 | 46 | Female | Han | Left | No | 1.6 | T1N0M0 | IDC-2 | Pos | Pos | Pos | No |
| Case_2 | 40 | Female | Han | Left | No | 4.1 | TisN0M0 | DCIS | Pos | Pos | Neg | No |
| Case_3 | 74 | Female | Han | Right | Yes | 7 | T4N0M0 | IDC-3 | NA | NA | NA | NA |
| Case_4 | 55 | Female | Han | Right | No | 3.9 | T2N0M0 | IDC-2 | Pos | Pos | Neg | No |
| Case_5 | 40 | Female | Han | Right | No | 3.1 | T2N1M0 | IDC-2 | Pos | Pos | Pos | No |
| Case_6 | 16 | Female | Han | Right | No | 1.8 | TisN0M0 | Papilla carcinoma | Pos | Pos | Neg | No |
| Case_7 | 54 | Female | Han | Right | No | 1.9 | T1N0M0 | IDC-2 | Pos | Pos | Neg | No |
| Case_8 | 38 | Female | Han | Left | No | 2.7 | T2N0M0 | IDC-2 | Pos | Pos | Neg | No |
| Case_9 | 49 | Female | Han | Left | No | 3 | T2N1M0 | IDC-2 | Pos | Pos | Neg | No |
| Case_10 | 57 | Female | Han | Left | Yes | 2.2 | T2N0M0 | IDC-2 | Pos | Pos | Neg | No |
| Case_11 | 45 | Female | Han | Right | Yes | 3.3 | T2N0M0 | IDC-3 | Neg | Neg | Neg | Yes |
| Case_12 | 53 | Female | Han | Right | Yes | 3.8 | T2N0M0 | IDC-3 | Pos | Neg | Pos | No |
| Case_13 | 31 | Female | Han | Left | Yes | 4.8 | T2N1M0 | IDC-2 | Neg | Neg | Pos | No |
| Case_14 | 32 | Female | Han | Left | No | 2.5 | T2N1M0 | IDC-2 | Neg | Pos | Pos | No |
| Case_15 | 58 | Female | Han | Left | No | NA | TXN0M0 | IDC-2 | Pos | Pos | Neg | No |
| Case_16 | 42 | Female | Han | Right | No | 2.6 | T2N1M1 | IDC-2 | Pos | Pos | Neg | No |
| Case_17 | 38 | Female | Han | Left | No | 5.1 | T3N1M0 | IDC-2 | Pos | Pos | Pos | No |
| Case_18 | 33 | Female | Han | Left | No | 10 | T3N2M0 | ILC | Pos | Neg | Neg | No |
| Case_19 | 52 | Female | Han | Right | No | 1.1 | T1N0M0 | IDC-2 | Pos | Pos | Neg | No |
| Case_20 | 61 | Female | Han | Right | No | 2.5 | TisN0M0 | Papilla carcinoma | Pos | Pos | Neg | No |
| Case_21 | 64 | Female | Han | Left | Yes | 1 | T1N1M0 | IDC-2 | Pos | Pos | Neg | No |
| Case_22 | 46 | Female | Han | Left | Yes | 3.6 | T2N1M0 | IDC-2 | Pos | Pos | Neg | No |
| Case_23 | 42 | Female | Han | Left | No | 3.3 | T2N0M0 | Mucinous carcinoma | Pos | Pos | Neg | No |
| Case_24 | 42 | Female | Han | Left | No | 2.5 | TisN0M0 | DCIS | Neg | Neg | Pos | No |
| Case_25 | 53 | Female | Han | Left | No | 2.3 | TisN0M0 | DCIS | Pos | Neg | Neg | No |
| Case_26 | 34 | Female | Han | Left | Yes | 4.4 | T2N0M0 | IDC-2 | Pos | Pos | Neg | No |
| Case_27 | 54 | Female | Han | Right | No | 2 | TisN0M0 | DCIS | Pos | Neg | Pos | No |
| Case_28 | 60 | Female | Han | Right | Yes | 1.1 | T1N1M0 | IDC-2 | Pos | Pos | Neg | No |
| Case_29 | 61 | Female | Han | Left | No | 1.5 | T1N2M0 | IDC-3 | Pos | Neg | Neg | No |
| Case_30 | 34 | Female | Han | Bilateral | Yes | 1.5 | NA | Papilla carcinoma | NA | NA | NA | NA |
| Case_31 | 43 | Female | Han | Left | Yes | 2.1 | T2N0M0 | IDC-2 | Pos | Pos | Neg | No |
| Case_32 | 31 | Female | Han | Right | No | 2.2 | TisN0M0 | DCIS | Pos | Pos | Neg | No |
| Case_33 | 50 | Female | Han | Left | No | NA | T1N2M0 | Occult carcinoma | NA | NA | NA | NA |
| Case_34 | 59 | Female | Han | Left | No | 3 | TisN0M0 | LCIS | Pos | Neg | Neg | No |
| Case_35 | 58 | Female | Han | Left | No | 1.6 | T1N2M0 | IDC-2 | Pos | Neg | Pos | No |
| Case_36 | 48 | Female | Han | Right | No | 5.1 | T2N0M0 | IDC-2 | Pos | Pos | Neg | No |
| Case_37 | 36 | Female | Han | Right | No | 2.4 | T2N1M0 | IDC-3 | Neg | Neg | Neg | Yes |
| Case_38 | 40 | Female | Han | Right | No | 1.4 | T1N1M0 | IDC-2 | Pos | Pos | Neg | No |
| Case_39 | 43 | Female | Han | Right | Yes | 3.4 | T2N0M0 | IDC | NA | NA | NA | NA |
| Case_40 | 40 | Female | Han | Left | Yes | NA | T2N1M0 | IDC-2 | Pos | Pos | Pos | No |
| Case_41 | 63 | Female | Han | Left | No | 0.5 | TisN0M0 | DCIS | Neg | Neg | Pos | No |
| Case_42 | 38 | Female | Han | Right | No | 2.4 | T2N1M0 | Micropapillary carcinoma | Pos | Pos | Neg | No |
| Case_43 | 61 | Female | Han | Left | Yes | 1.2 | T1N1M0 | IDC-2 | Pos | Pos | Neg | No |
| Case_44 | 35 | Female | Han | Right | No | 1.9 | T1N1M0 | IDC-2 | Pos | Pos | Neg | No |
| Case_45 | 38 | Female | Han | Right | Yes | 4.6 | T2N0M0 | DCIS | Pos | Pos | Neg | No |
| Case_46 | 59 | Female | Han | Left | Yes | 1.6 | T1N0M0 | IDC-3 | Neg | Neg | Neg | Yes |
| Case_47 | 24 | Female | Han | Left | No | NA | TXN0M0 | IDC-2 | Pos | Pos | Neg | No |
| Case_48 | 31 | Female | Han | Left | No | 5 | T3N0M0 | IDC-3 | Neg | Neg | Neg | Yes |
| Case_49 | 60 | Female | Han | Right | No | 2.7 | T2N1M0 | IDC-2 | Pos | Pos | Neg | No |
| Case_50 | 32 | Female | Han | Left | No | NA | T1N0M0 | IDC-2 | Pos | Pos | Neg | No |

BCOC, breast cancer and ovarian cancer; ER, estrogen receptor; PR, progesterone receptor; TNBC, triple-negative breast cancer; IDC, invasive ductal carcinoma; DCIS, ductal carcinoma *in situ*; Pos, positive; Neg, negative; NA, not available.

### Supplementary Table 2. Individual samples included in each mixed sample

| **Mixed sample** | **Case ID** |
| --- | --- |
| R10 | Case_1~ Case_10 |
| R20 | Case_1~ Case_20 |
| R30 | Case_1~ Case_30 |
| R40 | Case_1~ Case_40 |
| R50 | Case_1~ Case_50 |

### Supplementary Table 3. Twelve breast cancer susceptibility genes reported in the literatures^1,2^.

| GeneName | Refence Sequence |
| --- | --- |
| ATM | NM_000051 |
| BRCA1 | NM_007294 |
| BRCA2 | NM_000059 |
| CHEK2 | NM_007194 |
| PALB2 | NM_024675 |
| BARD1 | NM_000465 |
| RAD51C | NM_058216 |
| RAD51D | NM_002878 |
| TP53 | NM_000546 |
| CDH1 | NM_004360 |
| PTEN | NM_000314 |
| NF1 | NM_000267 |

1.	Dorling L, Carvalho S, Allen J, et al. Breast Cancer Risk Genes — Association Analysis in More than 113,000 Women. *New England Journal of Medicine.* 2021;384(5):428-439.

2.	Hu C, Hart SN, Gnanaolivu R, et al. A Population-Based Study of Genes Previously Implicated in Breast Cancer. *New England Journal of Medicine.* 2021;384(5):440-451.

### Supplementary Table 4. Pathogenic variants of breast cancer susceptibility genes detected in mixed DNA samples (P10-P20) by 550-panel sequencing.

| **Mixed**  **sample** | **BRCA1** | **BRCA2** | **PALB2** | **ATM** | **CHEK2** | **BARD1** | **RAD51C** | **RAD51D** | **TP53** | **CDH1** | **PTEN** | **NF1** |
| --- | --- | --- | --- | --- | --- | --- | --- | --- | --- | --- | --- | --- |
| P10 | - | - | c. 2488G>T | - | c.417 C>A | - | - | - | - | - | - | - |
| P20 | - | c.6443_6444insTA | c. 2488G>T | - | c.417 C>A | - | - | - | - | - | - | - |

### Supplementary Table 5. To identify which case carry the pathogenic variant found in the mixed DNA samples (P10 and P20) by site-specific Sanger sequencing

| **Mixed DNA sample** | **Individual DNA sample** | **PALB2:** c.2488G>T | **CHEK2:** c.417 C>A | **BRCA2:** c.6443_6444insTA |
| --- | --- | --- | --- | --- |
| P10 | Patient_1 | ╳ | ╳ | **-** |
|  | Patient_2 | ○ | ○ | **-** |
|  | Patient_3 | ╳ | ╳ | **-** |
|  | Patient_4 | ╳ | ╳ | **-** |
|  | Patient_5 | ╳ | ╳ | **-** |
|  | Patient_6 | ╳ | ╳ | **-** |
|  | Patient_7 | ╳ | ╳ | **-** |
|  | Patient_8 | ╳ | ╳ | **-** |
|  | Patient_9 | ╳ | ╳ | **-** |
|  | Patient_10 | ╳ | ╳ | **-** |
| P20 | Patient_1 | ╳ | ╳ | ╳ |
|  | Patient_2 | ○ | ○ | ╳ |
|  | Patient_3 | ╳ | ╳ | ╳ |
|  | Patient_4 | ╳ | ╳ | ╳ |
|  | Patient_5 | ╳ | ╳ | ╳ |
|  | Patient_6 | ╳ | ╳ | ╳ |
|  | Patient_7 | ╳ | ╳ | ╳ |
|  | Patient_8 | ╳ | ╳ | ╳ |
|  | Patient_9 | ╳ | ╳ | ╳ |
|  | Patient_10 | ╳ | ╳ | ╳ |
|  | Patient_11 | ╳ | ╳ | ╳ |
|  | Patient_12 | ╳ | ╳ | ○ |
|  | Patient_13 | ╳ | ╳ | ╳ |
|  | Patient_14 | ╳ | ╳ | ╳ |
|  | Patient_15 | ╳ | ╳ | ╳ |
|  | Patient_16 | ╳ | ╳ | ╳ |
|  | Patient_17 | ╳ | ╳ | ╳ |
|  | Patient_18 | ╳ | ╳ | ╳ |
|  | Patient_19 | ╳ | ╳ | ╳ |
|  | Patient_20 | ╳ | ╳ | ╳ |

Notes: “○” indicates the variant was detected in the corresponding cases; “╳” indicates the variant was not detected in the corresponding cases.

### Supplementary Table 6. Variants of uncertain significance in breast cancer susceptibility genes detected in mixed DNA samples (S10-S20) by 550-panel sequencing.

| **Mixed sample** | **BRCA1** | **BRCA2** | **PALB2** | **ATM** | **CHEK2** | **BARD1** | **RAD51C** | **RAD51D** | **TP53** | **CDH1** | **PTEN** | **NF1** |
| --- | --- | --- | --- | --- | --- | --- | --- | --- | --- | --- | --- | --- |
| S10 | c.A2481C | c.A4599C | - | - | - | - | - | c.G196A | - | - | - | - |
| S20 | c.A2481C | c.A4599C | - | - | - | c.T2277G | c.G424T | c.G196A | - | - | - | - |

### Supplementary Table 7. To identify which case carry the pathogenic variant found in the mixed DNA samples (S10 and S20) by site-specific Sanger sequencing

| **Mixed DNA sample** | **Individual DNA sample** | **BRCA1:**  **c.2481A>C** | **BRCA2: c.4599A>C** | **RAD51D: c.196G>A** | **BARD1: c.2277T >G** | **RAD51C: c.424G>T** |
| --- | --- | --- | --- | --- | --- | --- |
| S10 | Healthy_1 | ╳ | ╳ | ╳ | **-** | **-** |
|  | Healthy_2 | ○ | ╳ | ╳ | **-** | **-** |
|  | Healthy_3 | ╳ | ╳ | ╳ | **-** | **-** |
|  | Healthy_4 | ╳ | ╳ | ╳ | **-** | **-** |
|  | Healthy_5 | ╳ | ○ | ╳ | **-** | **-** |
|  | Healthy_6 | ╳ | ╳ | ○ | **-** | **-** |
|  | Healthy_7 | ╳ | ╳ | ╳ | **-** | **-** |
|  | Healthy_8 | ╳ | ╳ | ╳ | **-** | **-** |
|  | Healthy_9 | ╳ | ╳ | ╳ | **-** | **-** |
|  | Healthy_10 | ╳ | ╳ | ╳ | **-** | **-** |
| S20 | Healthy_1 | ╳ | ╳ | ╳ | ╳ | ╳ |
|  | Healthy_2 | ○ | ╳ | ╳ | ╳ | ╳ |
|  | Healthy_3 | ╳ | ╳ | ╳ | ╳ | ╳ |
|  | Healthy_4 | ╳ | ╳ | ╳ | ╳ | ╳ |
|  | Healthy_5 | ╳ | ○ | ╳ | ╳ | ╳ |
|  | Healthy_6 | ╳ | ╳ | ○ | ╳ | ╳ |
|  | Healthy_7 | ╳ | ╳ | ╳ | ╳ | ╳ |
|  | Healthy_8 | ╳ | ╳ | ╳ | ╳ | ╳ |
|  | Healthy_9 | ╳ | ╳ | ╳ | ╳ | ╳ |
|  | Healthy_10 | ╳ | ╳ | ╳ | ╳ | ╳ |
|  | Healthy_11 | ╳ | ╳ | ╳ | ╳ | ╳ |
|  | Healthy_12 | ╳ | ╳ | ╳ | ╳ | ╳ |
|  | Healthy_13 | ╳ | ╳ | ╳ | ╳ | ╳ |
|  | Healthy_14 | ╳ | ╳ | ╳ | ╳ | ╳ |
|  | Healthy_15 | ╳ | ╳ | ╳ | ╳ | ╳ |
|  | Healthy_16 | ╳ | ╳ | ╳ | ╳ | ╳ |
|  | Healthy_17 | ╳ | ╳ | ╳ | ╳ | ╳ |
|  | Healthy_18 | ╳ | ╳ | ╳ | ╳ | ╳ |
|  | Healthy_19 | ╳ | ╳ | ╳ | ╳ | ╳ |
|  | Healthy_20 | ╳ | ╳ | ╳ | ○ | ○ |

Notes: “○” indicates the variant was detected in the corresponding cases; “╳” indicates the variant was not detected in the corresponding cases.
